# Genome streamlining in CPR bacteria transitioning from soil to groundwater

**DOI:** 10.1101/2023.05.10.540099

**Authors:** Narendrakumar M. Chaudhari, Olga M. Pérez-Carrascal, Will A. Overholt, Kai U. Totsche, Kirsten Küsel

**Affiliations:** Aquatic Geomicrobiology, Institute of Biodiversity, Friedrich Schiller University, Jena, Germany; German Center for Integrative Biodiversity Research (iDiv) Halle-Jena-Leipzig, Leipzig, Germany; Hydrogeology, Institute of Geowissenschaften, Friedrich-Schiller-Universität Jena, Burgweg 11, 07749 Jena, Germany

## Abstract

To better understand the influence of habitat on the genetic content of Candidate Phyla Radiation (CPR) bacteria, we studied the effects of transitioning from soil to groundwater on genomic divergences of these organisms. Bacterial metagenome-assembled genomes (318 total, 35 of CPR bacteria) were generated from seepage waters and compared directly to groundwater counterparts. Seepage water CPR bacteria exhibited 1.24-fold greater mean genome size, while their inferred mean replication rate was 21.1% lower than groundwater lineages. While exploring gene loss and adaptive gains in closely related lineages in groundwater, we identified a zinc transporter, a surface protein, and a lipogylcopeptide resistance gene unique to a seepage Parcubacterium. A nitrite reductase gene unique to the groundwater Parcubacterium, likely acquired from pelagic microbes via horizontal gene transfer, was also identified. Groundwater Parcubacteria harbored nearly double the fraction (9.4%) of pseudogenes than their seepage kin (4.9%), suggesting further genome streamlining.

## Background

Genome size is a function of expansion and contraction, by gain and loss of DNA. Prokaryotes are susceptible to rapid losses and highly variable fluctuations in genetic content, oftentimes induced by selective environmental pressures^1,2^. Genome reduction leads to simplified metabolisms and lowered energetic requirements for cell duplication^3,4^. In bacteria, genome size is also habitat-dependent^5,6^. A global survey of genome size distribution suggested that aquatic bacteria harbor smaller genomes than their terrestrial counterparts, as the spatially and temporally diverse soil environment likely favors a broader genomic repertoire^7^. Aquatic habitats are often dominated by particularly tiny microbes adapted to oligotrophic conditions, as their high surface-to-volume ratios and superior transport systems render competitive advantages^3^. Such is certainly the case for free-living bacteria of the marine SAR11 lineage^8^, some marine Actinobacteria^9^, and freshwater Betaproteobacteria^10^.

Many groundwater microbiomes are dominated by taxa belonging to the Candidate Phyla Radiation (CPR), a large evolutionary radiation of bacterial lineages characterized by tiny cell sizes (0.1 to 0.7 µm) and compact genomes (0.3 to 1 Mbp)^11–17^. The genome size range of CPR organisms coincides with that of obligate symbiont bacteria^14^ and membrane-associated intracellular parasites^18^, which suggests a symbiotic lifestyle. Microscopic evidence points to an episymbiotic lifestyle for some CPR bacteria^15,19^, and the very few cultured representatives of this radiation, *e.g.*, Saccharibacteria (TM7), have been isolated solely as human symbionts^20,21^.

Given the lack of CPR isolates, most of what is known about these organisms’ genome sizes, metabolic potential, and lifestyles derives from metagenomic environmental surveys. In rhizosphere grassland soils, mean genome sizes were 0.61 ± 0.14 Mbp for Parcubacteria, 0.57 ± 0.11 Mbp for Saccharibacteria, and 0.79 ± 0.12 Mbp for Doudnabacteria^22^, while in Amazon grassland soils mean genome sizes were 0.5 ± 0.08 Mbp for Parcubacteria and 1.1 ± 0.2 Mbp for Microgenomates^23^. Unfortunately, from these limited genome size data alone, very little can be inferred about whether and to what extent CPR bacteria alter genome content in response to their habitat.

Encountering evolutionarily related yet ecologically differentiated microbes is rare^24,25^, as closely related microbes tend to segregate ecologically via accumulation of genetic changes and decreased genetic flow among them^26^. Studies on members of the *Methylophilaceae* family reported increased genome streamlining in cells collected from oligotrophic freshwaters compared to those colonizing sediments^24^. These observations were based on comparative whole-genome sequencing of bacterial isolates derived from habitats lacking representative CPR isolates. Given the paucity of CPR bacteria in soil^16,27^, even deep sequencing may fail to resolve high-quality genomes related to those detected in groundwater.

CPR bacteria are readily and preferentially mobilized from soils and vertically transported via seepage^16,27^ and potentially reach the underlying groundwater. Given the significantly greater abundance of CPR taxa in seepage than soil^16,27^, we attempted to detect soil-derived CPR bacterial genomes in seepage that were related to, or even the source of, CPR taxa in groundwater. Seepage waters were collected from soils of the preferential surface-recharge area and underlying vadose zone of the Hainich Critical Zone Exploratory (CZE). Experiments were conducted to discern the extent to which near-surface and groundwater microbes^17,28^ were genomically divergent, and whether taxonomic and/or genomic differences were due to gene loss and/or adaptive gain in response to habitat transition. Generally speaking, seepage CPR bacteria harbored larger genomes and exhibited lower replication rates than their groundwater counterparts. A summary of the findings of this study, which suggest that CPR bacteria undergo extensive genome diversification upon transitioning from near-surface soils to oligotrophic groundwater habitats, ensues.

## Results

### Elevated abundance of CPR bacteria in soil and vadose zone seepage communities

Twelve seepage water samples (six soil, six vadose zone) of the Hainich CZE were selected for metagenomic sequencing analysis, all of which were rich in CPR bacteria per 16S rRNA amplicon sequencing analyses. In these samples, abundances of *Cand.* Patescibacteria amplicon sequence variants (ASVs) ranged from 0.7 to 35% and 3.9 to 15% in soil and vadose zone seepage, respectively. Parcubacteria (up to 29% soil, 7.9% vadose zone) and Saccharimonadia (up to 2.9% soil, 9.3% vadose zone) were the most abundant CPR lineages detected, followed by Gracilibacteria (up to 1.8% soil, 0.3% vadose zone) and candidate division ABY1 (up to 0.8% soil, 0.4% vadose zone; Supp. Fig. 1A).

**Figure 1:**
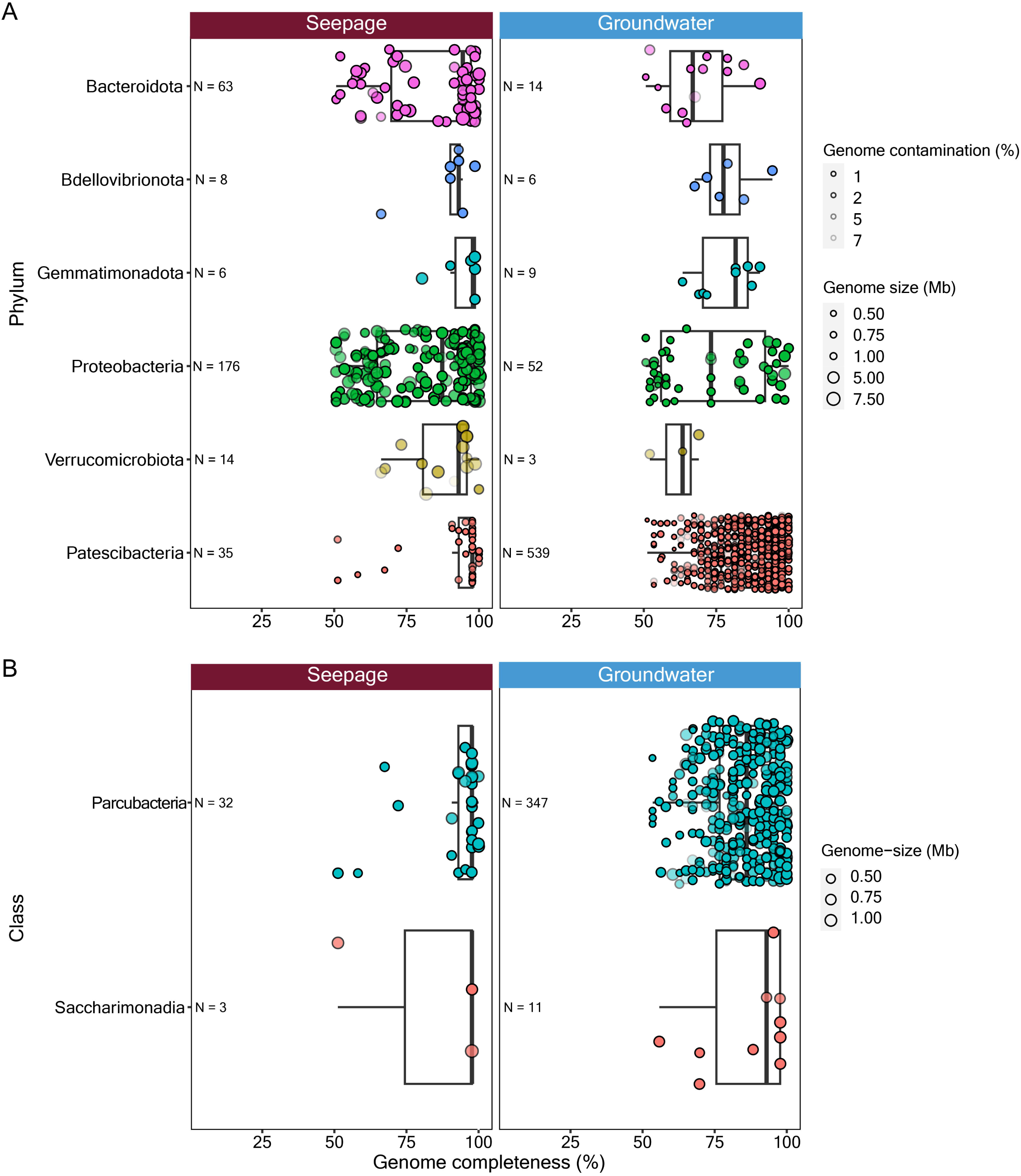
Summary of MAGs recovered from the seepage waters. **A**. Comparison of MAGs generated in this study to previously reported groundwater MAGs^17,28^. **B**. Comparison of MAGs representing each of two classes of Patescibacteria. Only phyla and classes represented by a minimum of three MAGs in both data sets were considered, in accordance with MIMAGs standards. Threshold of > 50% genome completeness and < 10% contamination applied to both plots. Circle size indicates genome size while shading intensity indicates extent of contamination.

The number of distinct CPR bacterial small subunit rRNA reads resulting from the sequenced metagenomes of these 12 samples exceeded the number of PCR amplicon derived ASVs by a factor of 2.5, with relative abundances ranging from 16.3 to 51.8% and 4.7 to 39.9% in soil and vadose zone seepage samples, respectively. Abundances of Parcubacteria, Saccharimonadia, candidate division ABY1, and Gracilibacteria reached up to 40, 3.5, 2.7, and 4.2% in the soil seepage communities, respectively, and 22.9, 11.1, 4.6, and 2.5%, in the vadose zone communities, respectively. Parcubacteria was the most abundant CPR bacterial lineage in both habitats.

Seepage water CPR bacterial abundances correlated strongly with previously reported measurements from the same ecosystem^16^. However, contrary to previous reports regarding non-CPR microbial community composition in oligotrophic groundwaters^17^, we found Proteobacteria to dominate both the soil (21.7 - 60.8%) and vadose zone seepage communities (32.7 - 40.8%), followed by Bacteroidota (3.2 - 13% and 6 - 34.6%, respectively), Verrucomicrobiota (3.2 - 7.4% and 1.2 - 9.2%, respectively), and Bdellovibrionota (0.6 - 2.3% and 0.9 - 5.7%, respectively; Supp. Fig. 1B).

### MAGs generated from soil and vadose zone seepage waters

Following appropriate binning and refinement, 318 non-redundant microbial MAGs were generated from the 12 distinct metagenomic assemblies. Of these, 139 MAGs (5 CPR bacteria) were medium-quality drafts (completeness ≥ 50%, < 90%; contamination < 10%) and 179 (30 CPR bacteria) were high-quality drafts (completeness ≥ 90%; contamination < 5%), as per Minimum Information about a Metagenome-Assembled Genome (MIMAG) standards^29^. With respect to genome completeness and contamination, the quality of seepage-associated MAGs representing bacterial phyla (Fig. 1A), including CPR clades (Fig. 1B), exceeded that of previously published data pertaining to underlying groundwater^17,28^. Both medium and high-quality MAGs were generated from both the soil and vadose zone seepage water samples.

Of 35 resulting CPR bacterial MAGs, 32 represented Parcubacteria (mean genome size 203.4 Kbp ± 234.5) and three Saccharimonadia (mean genome size 1031.9 Kbp ± 200.3). We failed to generate high-quality MAGs from any other CPR lineages (*e.g*., Gracilibacteria, Microgenomatia, candidate division ABY1), likely due to the scarcity of representative nucleic acid sequences. Phylogenetic reconstruction based on 71 conserved gene-product sequences of all MAGs generated from seepage waters confirmed the CPR as a monophyletic group composed of two lineages: Parcubacteria and Saccharimonadia (Supp. Fig. 2).

**Figure 2:**
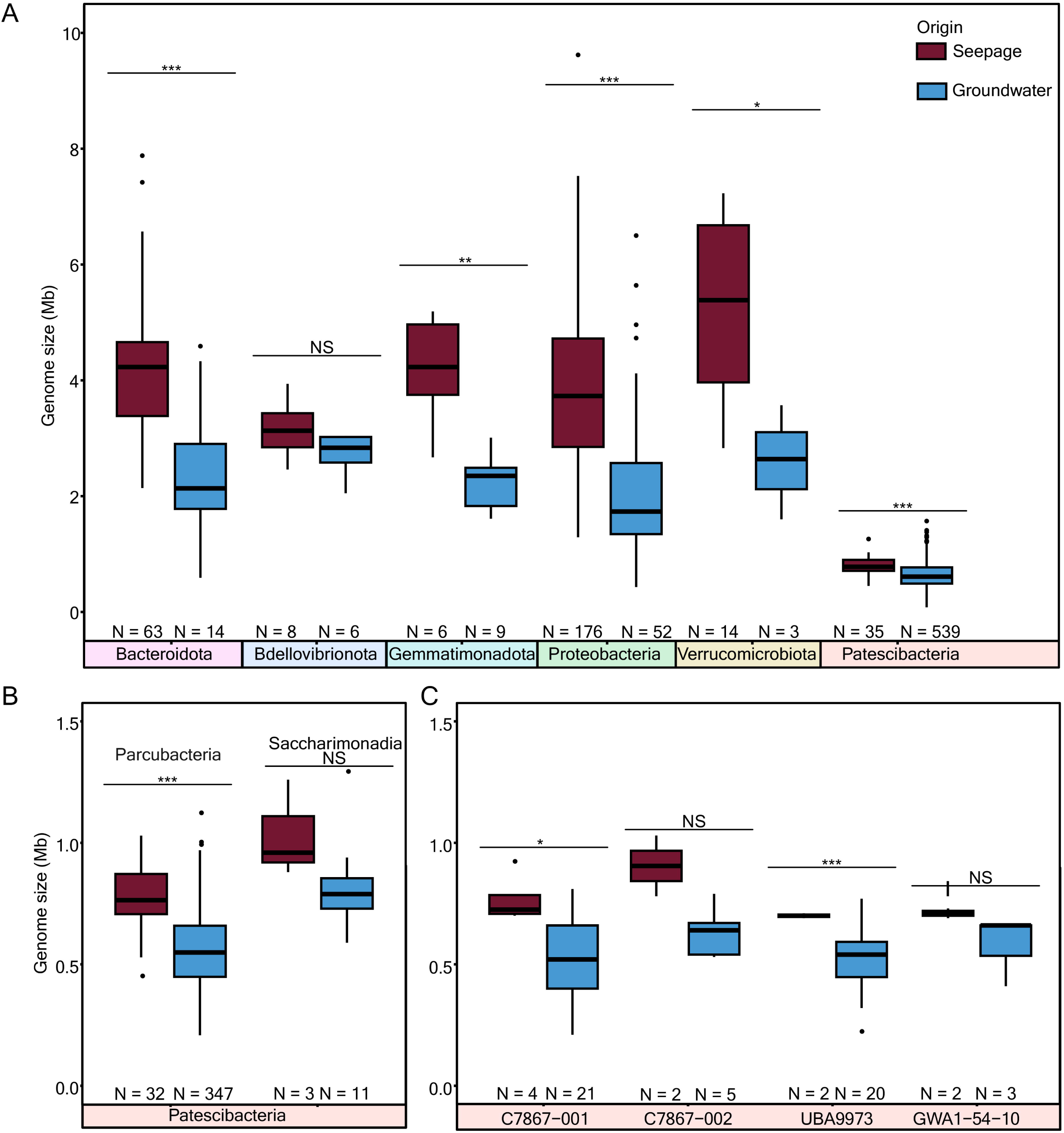
Genome size differences in seepage and groundwater bacteria. **A**. major bacterial phyla, **B**. Patescibacteria classes, and **C**. genera within the Parcubacteria class of Patescibacteria. Statistical significance is denoted as not significant (NS) for P > 0.05, * for P ≤ 0.05, ** for P ≤ 0.01, and *** for P ≤ 0.001.

### Larger bacterial genomes in seepage waters than groundwaters

Representatives of most major bacterial phyla, including Bacteroidota, Gemmatimonadota, Proteobacteria, Verrucomicrobiota, and the CPR harbored significantly larger genomes in seepage communities than their counterparts in groundwater communities (Fig. 2A). High-quality Parcubacterial MAGs were significantly larger (25%, p = 1.47 × 10^-6^) in seepage water samples (0.802 Mbp) than groundwater samples (0.641 Mbp). Seepage water borne MAGs representative of the Parcubacterial genus C7867-001 were significantly larger than MAGs representing the same genus in groundwaters (p = 0.019; Fig. 2C). Mean genomic GC content was 1.1-fold greater in seepage borne bacteria than their groundwater relatives.

### Groundwater borne bacteria replicate faster than their seepage water relatives

Microbial replication rates fluctuate in response to environmental conditions^30^. Studying closely related bacterial groups from dissimilar habitats, *e.g*., seepage vs. groundwater, facilitates important observations and empirical inferences pertaining to the impact of various ecological stimuli on proliferation. We found that estimated mean *in-situ* growth rate indices (GRiD) derived from groundwater borne MAGs were significantly greater than those of seepage water borne MAGs of Acidobacteriota (1.23-fold, p = 1.06 × 10^-2^), Bacteroidota (1.2-fold, p = 1.98 × 10^-6^), Gemmatimonadota (1.36-fold, p = 8.42× 10^-10^), Verrucomicrobiota (1.17-fold, p = 4.28× 10^-2^), and Patescibacteria (1.27-fold, p = 6.35× 10^-28^; Fig. 3A). Differences in replication rate were particularly pronounced across classes and genera of CPR bacteria (Fig. 3B-C). As a general trend, seepage water microbes replicated at lower rates than their groundwater counterparts.

**Figure 3:**
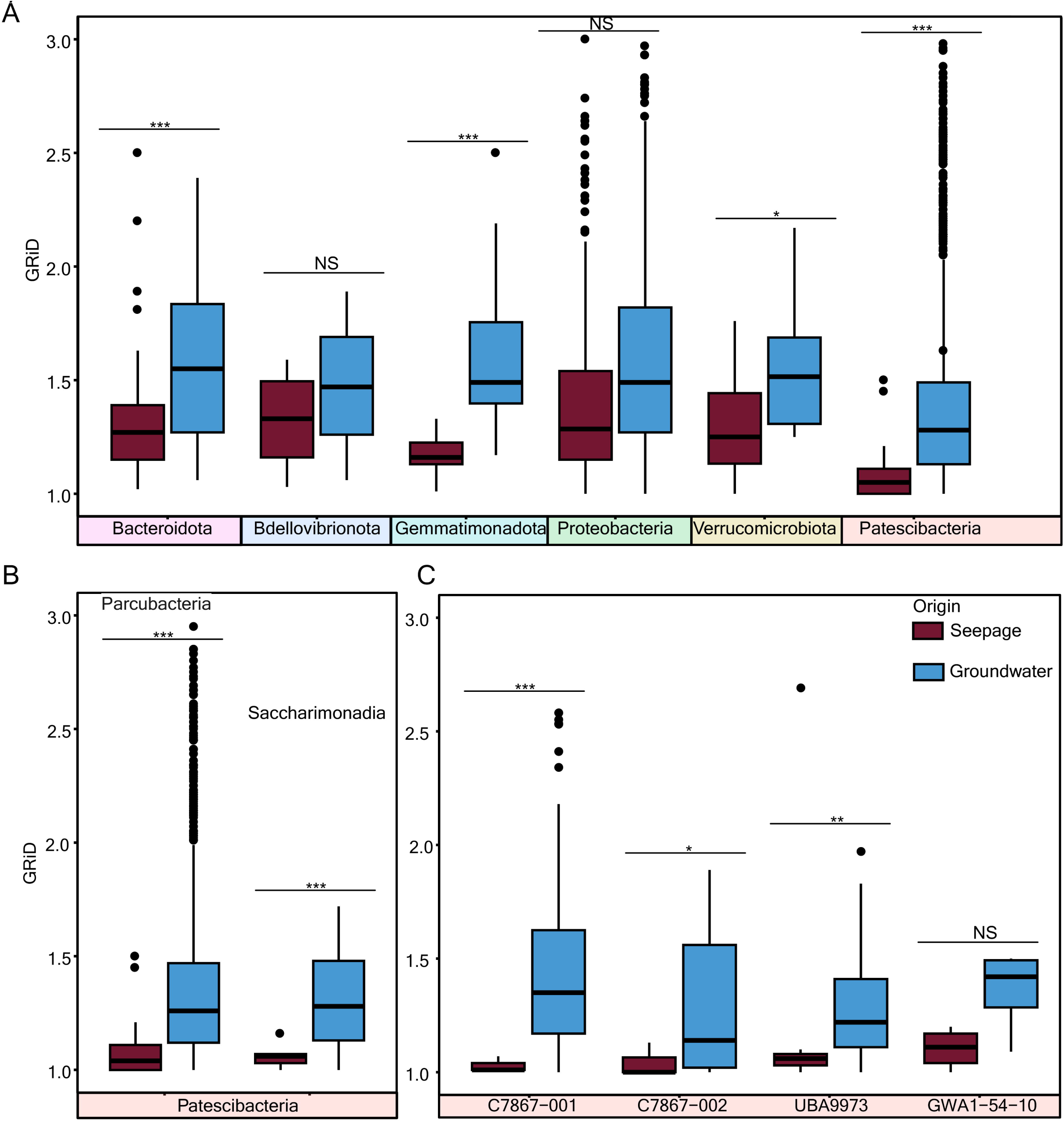
Estimated growth rate index (GRiD) distribution of MAGs of all bacterial phyla (A), CPR bacterial classes (B), and CPR bacterial genera (C). Statistical significance was calculated using the t-test function with false discovery rate (FDR) correction in R (v.4.2.2), denoted here as not significant (NS) for P > 0.05, * for P ≤ 0.05, ** for P ≤ 0.01, and *** for P ≤ 0.001.

### Habitat-distinct traits in closely related Parcubacteria

Studying genomically similar microorganisms in different environments using only metagenomics is challenging, especially when the abundance of such targets is particularly low in one or more of the habitats probed. A comparative genome analysis based on mean nucleotide and amino acid identity (AAI) conducted in parallel with phylogenomic placement (Supp. Fig. 2) of available MAGs revealed a pair of closely related MAGs belonging to the same genus of Parcubacteria – one from seepage and the other from groundwater. This facilitated exploration of genomic and functional differences that may drive the adaptation of microbes transitioning to new habitats.

A seepage-borne Parcubacterium MAG (ADI-DC-SW-Bin061) belonging to genus C7867-001 was nearly twice (697 Kbp) as large as that of a closely related groundwater-borne Parcubacterium MAG (H51-Bin103; 388 Kbp) of the same genus. However, estimated completion sizes were only 97% and 72%, respectively (Table 1). Mean AAI between these MAGs was 92.44%. Nearly half of all gene clusters identified in these Parcubacterial MAGs were unannotated against KEGG or COG, most of which were deemed hypothetical proteins. There were 400 gene clusters unique to the seepage Parcubacterium genome and 94 gene clusters unique to its groundwater counterpart (Fig. 4A). The 363 gene clusters shared by the two genomes accounted for 79% of the groundwater-borne MAG’s gene density.

**Figure 4:**
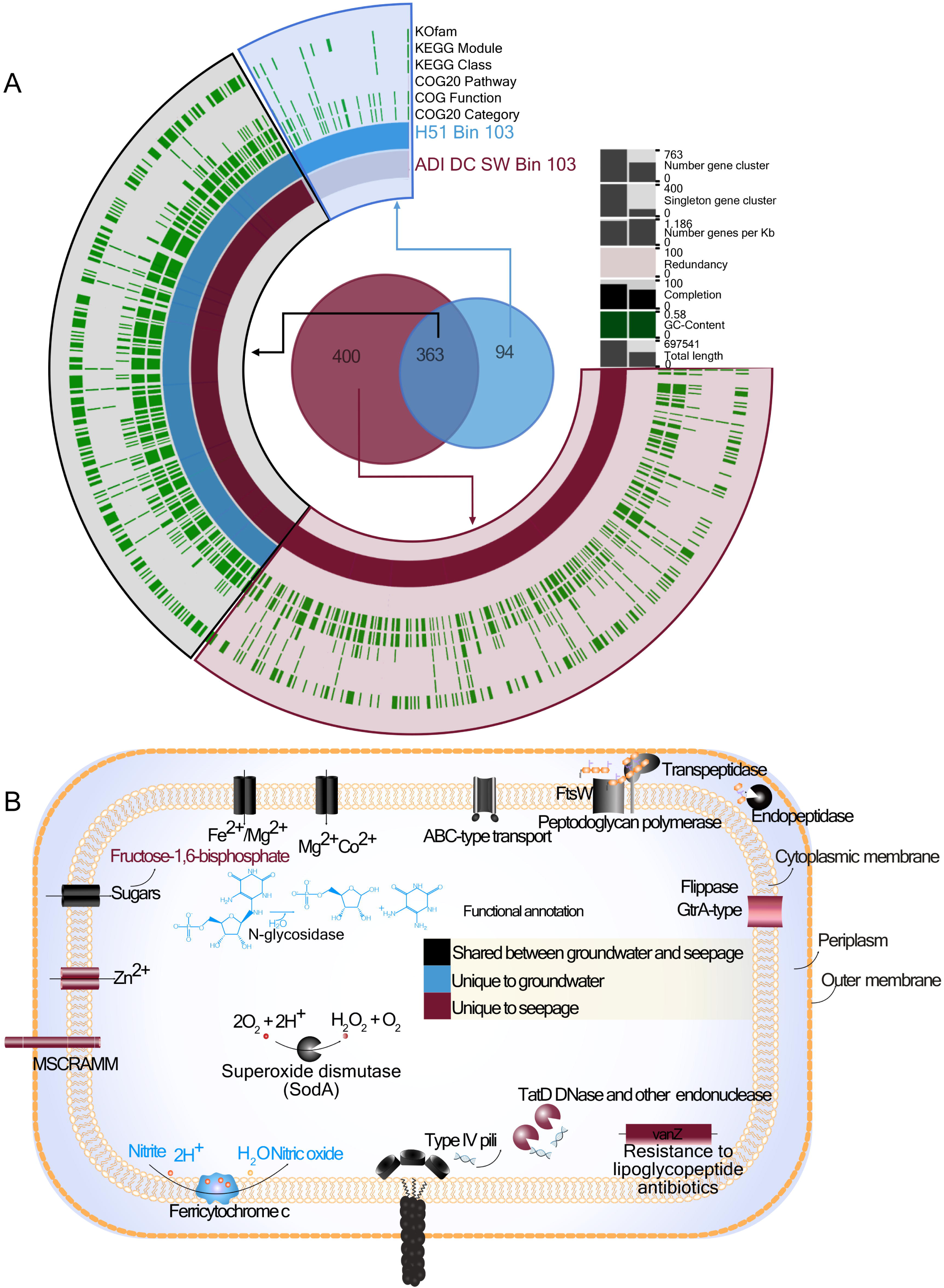
**A.** Functional profile of genome-specific and conserved gene clusters between a pair of Parcubacteria MAGs. Each bar along the radius of the map represents a distinct gene cluster. Genes from the groundwater MAG (sky blue) and seepage MAG (brown) are shown alongside respective functions (green) derived from COG and KEGG pathways. **B.** Illustration of genome-specific and shared cellular and metabolic features between the two Parcubacteria MAGs.

**Table 1:** Comparison of features of seepage (this study) vs. groundwater^17,28^ genomes of Parcubacterial genus C7867-001.

| Source | Seepage | Groundwater |
| --- | --- | --- |
| Genome name | ADI-DC-SW-Bin061 | H51-Bin103 |
| Genome size (Kbp) | 697.5 | 388.3 |
| No. of contigs | 2 | 50 |
| Coding genes | 775 | 438 |
| Pseudogenes | 38 | 41 |
| Percentage of pseudogenes | 4.9% | 9.4% |
| tRNAs | 43 | 25 |
| GC content | 57.9 | 58.2 |
| Genome completeness | 97% | 72% |
| Genome contamination | 0% | 0% |
| Fraction of read coverage within binned genomes in respective sample | 31.1% in vadose zone water sample<br>DC_SW_02_SPN_11032020 | 0.33% in<br>groundwater well<br>H51 |

A detailed screening of gene functions revealed the presence of both common and specific habitat-derived structural and metabolic features. A sugar transporter, ion transporters for iron and magnesium, an ABC-type transport system, and superoxide dismutase were encoded by both genomes. The seepage Parcubacterium encoded a zinc ion transporter, a flippase, a TatD nuclease family protein, and a surface protein from a family of microbial surface components that recognize adhesive matrix molecules (MSCRAMM, COG4932). Sugar utilization functions were more abundant in seepage-borne CPR bacteria, linking to a more heterotrophic, nutrient rich environment wherein opportunities to utilize sugar intermediaries as energy sources abound. Incomplete pathways for the metabolism of methane, vitamins, cofactors, and amino acids were also detected in the seepage-borne CPR bacterial genome.

The groundwater MAG encoded an enzyme that reduces nitrite to nitric oxide. While this enzyme is common to groundwater CPR bacteria^15,17^, it was not detected in seepage CPR bacteria (Fig. 4B). This is a unique example of functional and genomic divergence between two closely related CPR bacteria inhabiting different, yet interconnected, environments. In addition, a gene encoding N-glycosidase unique to the groundwater-borne CPR bacterial genome. This enzyme plays a role in the cleavage of N-glycosidic bonds of riboflavin intermediates^31^. While pathways involved in the biosynthesis of riboflavin have been reported in groundwater CPR bacteria^18^, aspects pertaining to its catabolism (*e.g.*, use of intermediates as an energy source in low nutrient environments) remain poorly understood.

### Genome streamlining in groundwater Parcubacteria

To determine whether differences in genome size resulted from habitat-dependent streamlining, we quantified pseudogenes and/or genes in the process of being lost or becoming non-functional. As was expected, the seepage-borne MAG harbored a smaller fraction of pseudogenes (4.9% of coding genes) than the groundwater MAG (9.4% of coding genes; Table 1). A general comparison of all CPR bacterial MAGs showed a similar trend, with groundwater-borne MAGs bearing significantly more pseudogenes than their seepage counterparts (Fig. 5A). This disparity is likely a consequence of genome reduction and loss of metabolic function in groundwater CPR bacteria, which might have rendered improved fitness to the oligotrophic conditions and symbiotic lifestyle.

**Figure 5:**
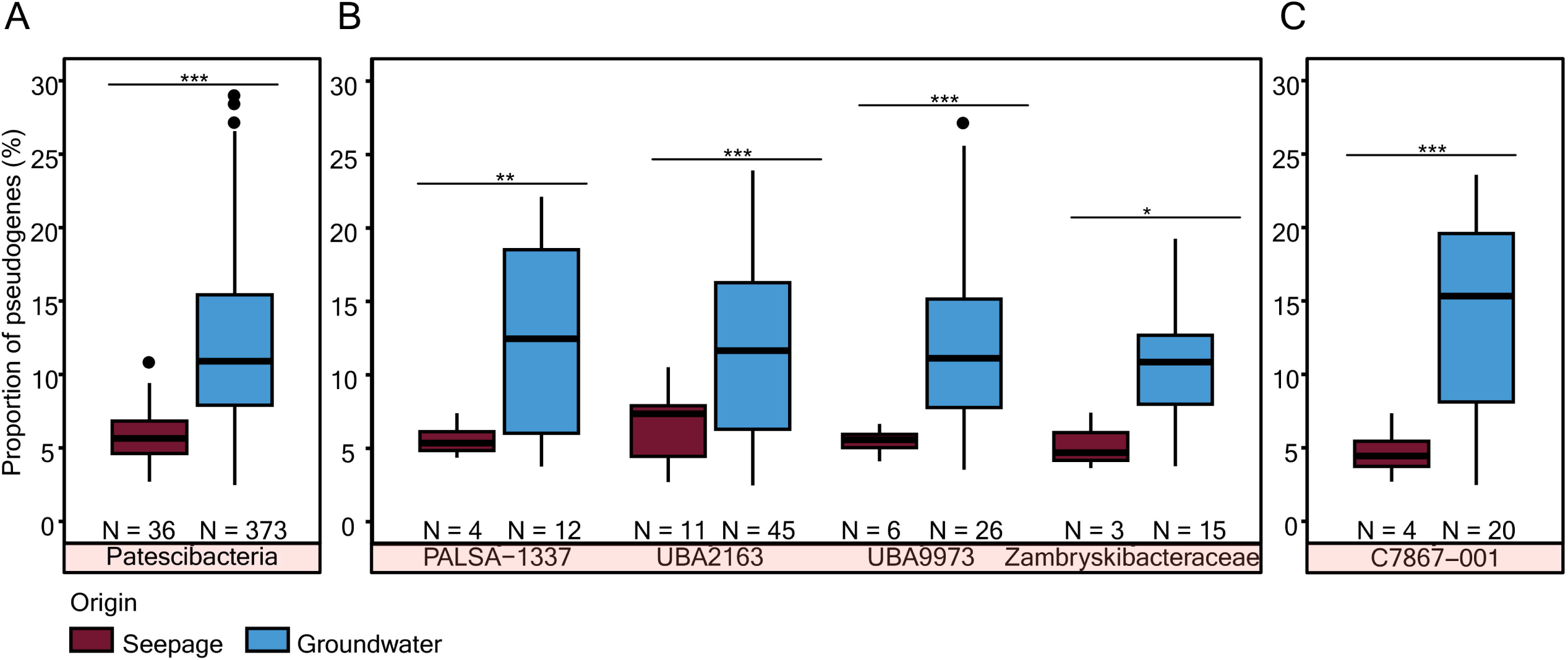
Comparison of pseudogene frequencies in MAGs derived from seepage and groundwater. The percent contribution of pseudogenes is compared for MAGs of CPR bacteria (A), families of Parcubacteria (B), and a genus of Parcubacteria (C) found both in seepage (this study) and groundwater^17,28^. Statistical significance is denoted by * for P ≤ 0.05, ** for P ≤ 0.01, and *** for P ≤ 0.001.

While low GC content is typically an indicator of limited nitrogen availability and genome streamlining^2,3,32^, we did not observe significant differences in the GC content of the seepage vs. groundwater MAGs (Supp. Fig. 3).

## Discussion

Our experimental approach afforded the ability to study microbes transitioning from soil to underlying groundwater habitats, a feat hitherto achieved only in pelagic waters and sediments^8,24,33^. By focusing on differing genomic characteristics between seepage and groundwater-borne microbes, we confirm the occurrence of closely related microbes bearing unique genetic content that supports contrasting lifestyle strategies in these interconnected yet distinct habitats.

Seepage water connects surface habitats like soil to the underlying unsaturated vadose zone and groundwater. Samples collected from both soil and vadose zone seepage water were dominated by Patescibacteria, with relative abundances reaching 50.4% and 41.1%, respectively, and Parcubacteria accounting for up to 40 and 22.9% of the microbial population, respectively. Remarkably, Patescibacteria accounted for a mere 0.55% of forest soil microbial relative abundance and Parcubacteria were up to three orders of magnitude more abundant in soil seepage, whereas members of Actinobacteria and Acidobacteria, which tend to adhere to soil matrices, were underrepresented in seepage waters^16^. Ergo, the transition of microbial communities from soil to groundwater appears to favor particular taxa, seemingly based on cellular attachment to soil matrices, surface charges, and/or other hitherto unresolved specific traits or lifestyle determinants.

We exploited this disparity in mobility behavior and generated 318 distinct bacterial MAGs from seepage waters. The availability of several seepage water associated MAGs, presumably originating from overlying soil, and 1,224 groundwater MAGs^28^ facilitated highly informative comparative analyses of bacterial genomes originating from different habitats. Within the same lineage, seepage bacteria tended to maintain larger genomes than groundwater denizens (Fig. 2A). This trend was prominent in Proteobacteria and the classes, and genera of Patescibacteria (Fig. 2B-C). Inter-habitat (seepage vs. groundwater) differences in CPR bacterial genomes were more pronounced than intra-habitat differences (various locations within a seepage area or about a groundwater transect). However, variations in the estimated genome quality of MAGs recovered from both habitats might have contributed to the observed genome size differences. Overall, our results suggest that the transition of CPR bacteria from complex, heterogeneous surface soil environments to more consistent and oligotrophic groundwaters is accompanied by a reduction in genome size, although many likely live episymbiotically^15^.

Closely related microbes are rarely encountered across different habitats^24^. Between two CPR (Parcubacteria) bacterial MAGs of the same genus (C7867-001, AAI = 92.44%), one retrieved from seepage and the other from groundwater, a greater number of genes were exclusive to the seepage-borne MAG than its groundwater counterpart. While our interpretations are somewhat limited by incomplete annotation, this could result from the shedding of unnecessary genes upon transitioning to oligotrophic groundwaters scarce in energy sources. Annotated genes unique to the seepage Parcubacteria encoded a zinc transporter, which might facilitate the selective uptake of zinc from surrounding soils, a surface protein (clfA) involved in attachment, and an antibiotic resistance protein that prevents binding of lipoglycopeptide antibiotics^34^. The nitrite reductase gene (*nirK*) was unique to groundwater Parcubacteria, likely in response to local exposures to nitrate and/or nitrite. The products of this gene might also contribute to denitrification processes in groundwater^13^. Type-IV pili, typically responsible for natural competence and extracellular DNA uptake^35^, were present in both Parcubacteria. However, only the seepage borne Parcubacterium carried the *tatD* gene with DNase, which could afford a means of utilizing extracellular DNA in soils^36^.

Genome streamlining is an adaptive strategy used by bacteria to save energy. Cells encountering deleterious and/or energy limited conditions shed genetic content and machineries whose upkeep is no longer energetically worthwhile. We observed greater fractions of suspected pseudogenes in the groundwater MAGs. Pseudogenes accounted for roughly 10 and 5% of the coding genes in the genome of the groundwater and seepage borne Parcubacteria, respectively. Most often, pseudogenes were functional in the past but underwent mutational changes resulting in their removal over the course of evolution^37^. The greater fraction of such genes in groundwater microbes, is indicative of a higher probability of gene loss and further genome streamlining.

Bacterial genome size is oftentimes correlated with genomic GC content, with compact genomes of obligate endosymbionts presenting the lowest GC contents^38^. Groundwater borne CPR bacterial MAGs of genus C7867-001 exhibited 8.9% lower (p=0.023) mean genomic GC contents than their seepage relatives. Given the higher production cost of guanine and cytosine and greater intracellular availability of adenine, tyrosine, and uridine^39,40^, low GC content is both energetically favorable and a bolster to fitness in oligotrophic environments.

Groundwater borne bacteria replicated faster than seepage bacteria. This trend held for all the classes, families, and genera of superphylum *Cand*. Patescibacteria. Faster replication rates can be linked to smaller genomes, which simply require less time and energy to replicate^41^. Groundwater borne CPR bacteria appear to thrive at this site, as their lifestyle affords them the ability to grow faster in this nutrient-poor habitat than their relatives in seepage habitats. Alternatively, groundwater might simply provide better opportunities for encountering host organisms, as elevated replication rates have been reported for CPR bacteria physically attached to host cells^15^. In our sequential filtration pipeline, attached CPR bacteria should have been retained on the larger 0.2 µm filters. Accordingly, groundwater borne MAGs of class Parcubacteria generated from 0.2 µm filter fractions had significantly greater GRiD values than those generated from 0.1 µm filter fractions. Co-occurrence patterns hinted at a few potential hosts in groundwater, such as members of phyla Nanoarchaeota, Bacteroidota, MBNT15, Bdellovibrionota, Nitrospirota, and Omnitrophota^17^. Thus, CPR bacteria might form transient attachments to hosts in groundwater to bolster replication for limited periods of time^17^.

Ultimately, the results of this investigation demonstrate that CPR bacteria, characterized by ultra-compact genomes and minimal biosynthetic and metabolic potential, further optimize their efficient lifestyle following mobilization from soil habitats. The observed gene loss of 11% and greater fraction of pseudogenes in groundwater borne CPR bacteria than their seepage counterparts both contribute significantly to genome streamlining. The presence of specific genes in one but not both closely related Parcubacteria exemplifies niche adaptation to either seepage water or groundwater.

## Methods and Materials

### Sample collection, DNA extraction, and metagenomic sequencing

Seepage samples were collected from soils and locations about the vadose zone at the Hainich Critical Zone Exploratory (CZE) in Thuringia, Germany. Six tension-controlled lysimeters installed at the topsoil/subsoil and subsoil/parent rock interfaces at roughly 30 cm to 60 cm below the soil surface were used to collect seepage samples. In addition, six collectors were used to sample the free drainage percolating through the vadose zone at depths ranging from 97 to 169 cm. All samples were collected between December 2019 and March 2020. Filtration of soil seepage water and vadose zone seepage water samples was accomplished using 0.1 and 0.2 µm membrane filters (Supor®, Pall), respectively. Filters were immediately stored at −80 °C. DNA extraction was carried out from the filters using a DNeasy® PowerSoil® kit in accordance with manufacturer’s protocols (Qiagen, USA). Shotgun metagenomic sequencing was carried out using an Illumina NovaSeq 6000 SP Reagent kit (v1.5; 300 cycles) on an Illumina NovaSeq6000 sequencing platform.

### Read quality filtering, metagenomic assembly, genome binning, and bin refinement

Metagenomic sequencing yielded an average of 78.8 ± 7.7 million reads per sample. Following quality filtering via the bbduk script (BBMap version 38.96)^42^, only high-quality reads were retained. Read error corrections were processed using bbnorm (BBMap)^42^ followed by *de novo* assembly using SPAdes (v3.13.0) (using --meta mode)^43^. Contigs longer than 1,000 bp were binned using maxbin2^44^, metabat2^45^, and binsanity^46^. Metawrap^47^ refinement (using filters -c 50 -x 10) was then carried out to refine bins obtained from the three binning algorithms and obtain the best representative MAGs. Bins (MAGs) were manually refined via visual inspection of contig coverage and sequence composition profiles, using the Anvi’o (v.7) suite^48^ to further improve the quality of refined genomes. A final genome-quality assessment was carried out using the CheckM^49^ workflow (v.1.1.3) with a lineage-specific set of marker genes for all CPR bacteria. Only MAGs having at least 50% genome completeness and at most 10% redundancy/contamination were retained for subsequent comparisons.

### Taxonomic annotation and phylogenetic analysis of MAGs

Taxonomic annotations of the MAGs selected for analysis were carried out with GTDB-Tk^50^ (v1.5.1) using GTDB (release 202) as a reference database^50^. A Maximum-likelihood (ML) phylogenetic tree was constructed based on concatenated alignments of the amino-acid sequences of 71 single-copy core genes found in all the MAGs using FastTree (v2.1.10) (1,000 bootstrap replicates)^51,52^.

### Estimation of growth rates

Metagenomic read coverages of origin of replication and terminus of the individual MAGs were used to calculate differences in genome copy number caused by ongoing replication. Using GRiD (v.1.3), this information was then used to estimate the growth rate index (GRiD)^53^ for each MAG having sufficient mean genome read coverage (at least 0.7)^53^. This index is proportional to the growth rates of organisms being analyzed^53^.

### Comparative genomics of CPR bacterial MAGs from near-surface and groundwater communities

The average amino-acid identities (AAI) of all possible pairs of CPR bacterial MAGs from seepage (*n* = 35) and groundwater (*n* = 584) were calculated using EzAAI (v.1.2.0)^54^. A pair of closely related CPR bacterial MAGs yielding an AAI value of 92.44% (the highest value observed) was selected for in-depth comparative analysis of genetic content and metabolic potential. One of these MAGs arose from a seepage sample, the other from a groundwater sample. Contigs from these Parcubacterial MAGs were processed with Prodigal^55^ (v.2.6.3) to identify open reading frames. All protein-coding genes from the two genomes were clustered using the Anvi’o pan-genome suite with default parameters. COG^56^ and KEGG^57^ functions were annotated using respective databases within Anvi’o. Reverse translated DNA amino acid sequences of the two Parcubacterial MAGs were mapped to the KEGG database using the bidirectional best blast-hits method within the KAAS web server^58^, and output was manually screened for individual genes specific to respective genomes. Pseudogenes were identified using Pseudofinder^59^ (v.1.1.0), and counts were normalized to percentage of total annotated genes to simplify direct comparison.

## Additional information

Correspondence and requests for materials should be addressed to Kirsten Küsel.

## Acknowledgments

NMC gratefully acknowledges the support of the German Centre for Integrative Biodiversity Research (iDiv) Halle-Jena-Leipzig funded by the Deutsche Forschungsgemeinschaft (DFG, German Research Foundation; FZT 118-230 202548816). OP-C gratefully acknowledges support from the DFG under Germany’s Excellence Strategy - EXC 2051 - Project-ID 390713860. This study is part of the Collaborative Research Centre AquaDiva of the Friedrich Schiller University Jena, funded by the DFG-SFB 228 1076-Project Number 218627073. The authors acknowledge seepage water sampling and further processing by K. Lehmann, D. Chaturangani, and F. Gutmann.

## Author contributions

KK, NMC, and WAO designed the study. NMC performed the bioinformatics and data analyses with help of WAO and OP-C. KUT designed and managed the field installations of seepage and groundwater monitoring. NMC and KK wrote the manuscript. All authors discussed the results and implications and commented on the manuscript at all stages. All authors have seen and approved the manuscript, and it hasn’t been accepted or published elsewhere.

**Supp. Figure 1: Microbial community composition of soil seepage and vadose zone seepage waters** based on the relative abundance of ASVs obtained from 16S rRNA amplicon sequencing (A) and the fraction of metagenomic reads mapped to the SSU rRNA gene (B).

**Supp. Figure 2:** Phylogenetic placement of MAGs generated in this study based on concatenated protein sequence alignments of a set of bacterial single-copy core genes (see Methods for more details). A groundwater Parcubacterium MAG (sky-blue branch within Parcubacteria clade) closely related to the seepage borne Parcubacterium MAG (brown branch within Parcubacteria clade) discussed throughout this manuscript is included.

**Supp. Figure 3:** Comparison of genomic GC contents of MAGs generated from seepage and groundwater samples for bacterial phyla (A), CPR classes (B), and genera within CPR class Parcubacteria (C).

