## Supplementary figures and images for "Genome streamlining in CPR bacteria transitioning from soil to groundwater"

### Supp. Figure 1

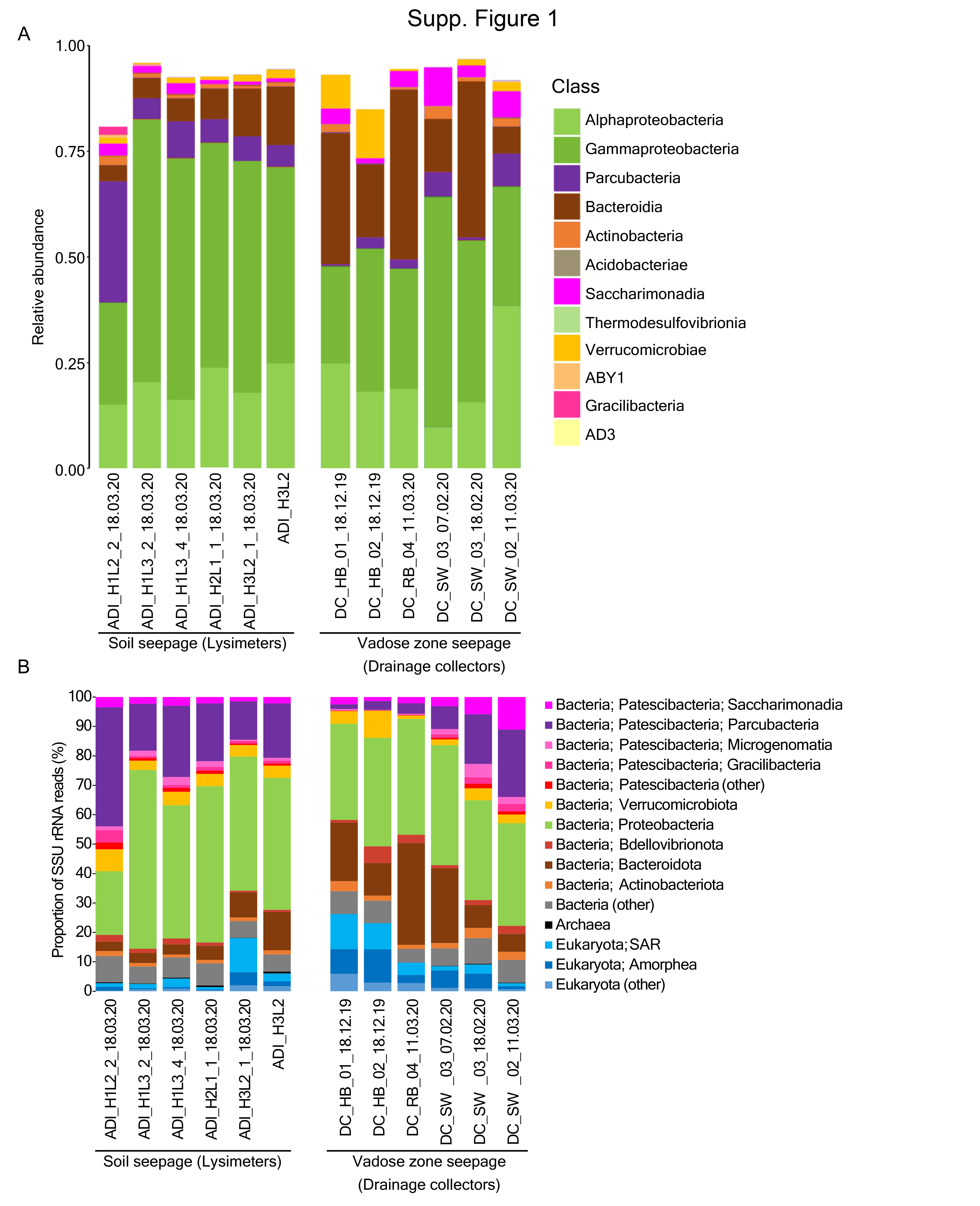

### Supp. Figure 2

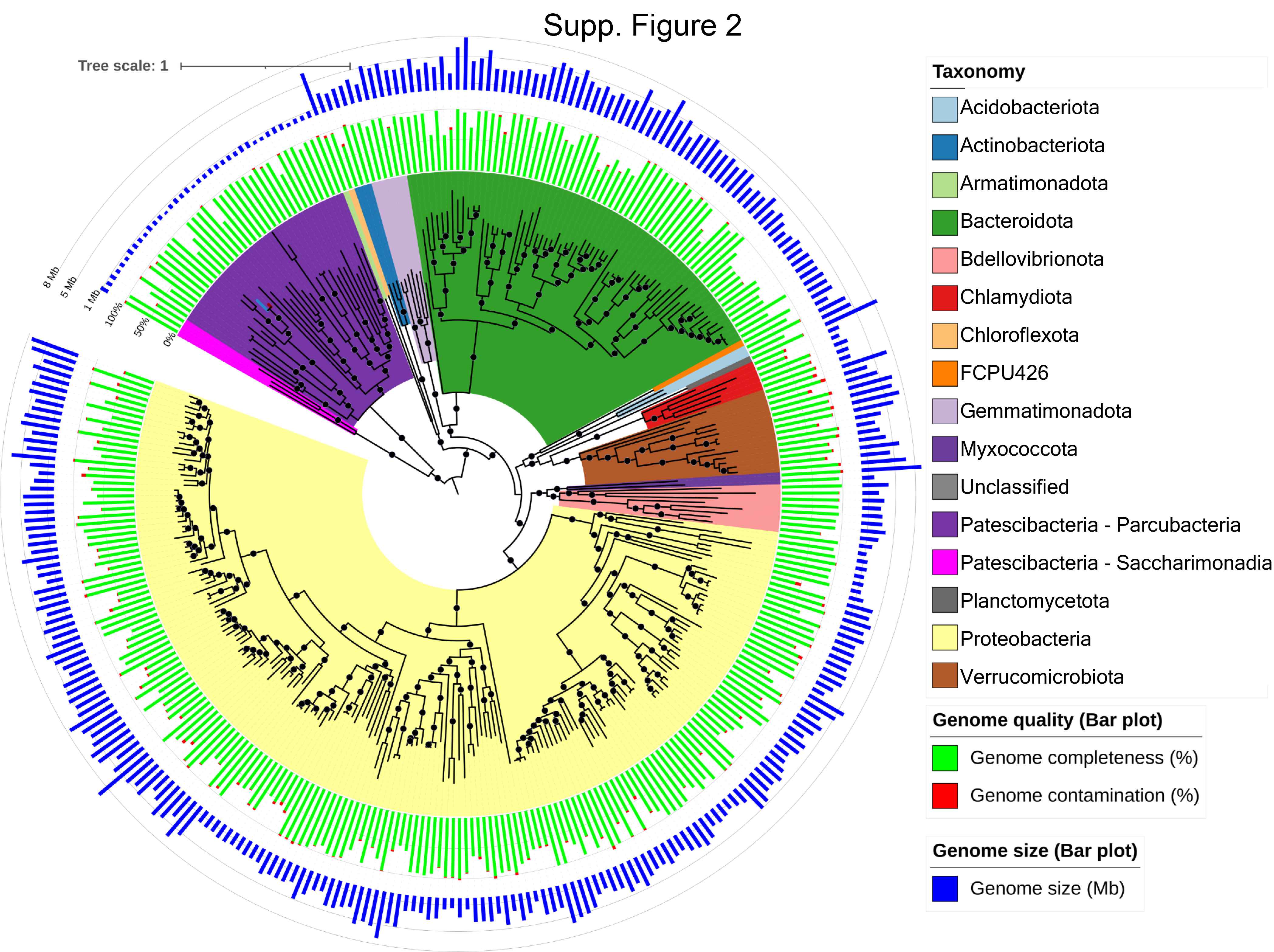

### Supp. Figure 3

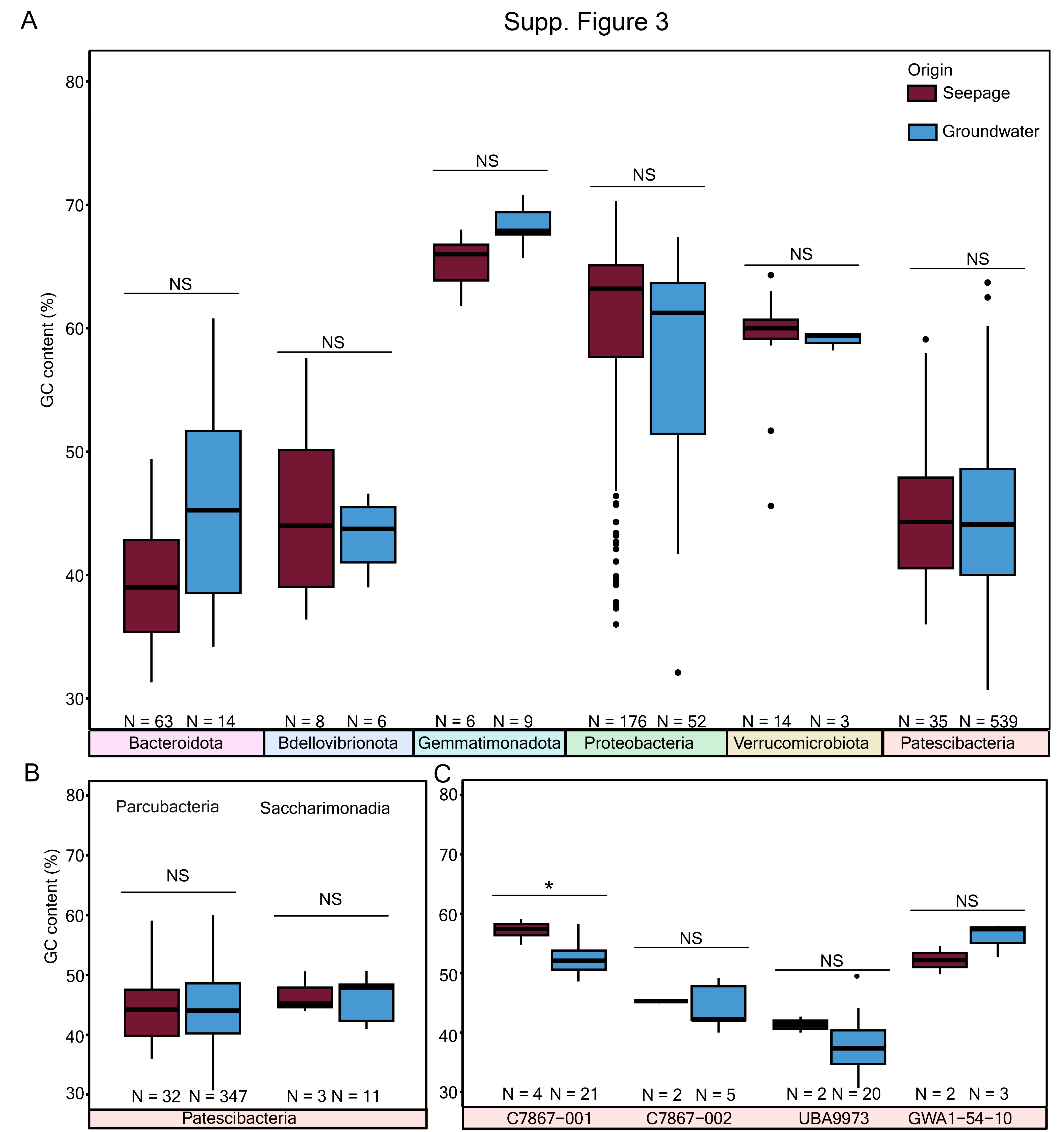
